# Dynamics of a mixed beech (*Fagus silvatica* L.) and hornbeam (*Carpinus betulus* L.) natural regeneration in North-East France: evolution of stand density and species composition with an insight into tree mortality factors

**DOI:** 10.1101/2025.02.28.640759

**Authors:** N. Le Goff, J.-M. Ottorini, F. Ningre

## Abstract

**Context:** Mixed beech and hornbeam regenerations are quite common in the east of France but their management is based more on general principles linked to ecosystem functions or services than on silvicultural practices. This is why it is so useful to establish stand trajectories to characterizing the evolution of density during stand development, as well as individual tree mortality models to predict how each species may react, particularly in a changing environment.

**Aims:** This study was conducted to establish size-density trajectories of mixed even-aged beech and hornbeam natural regenerations, using an already tested piecewise polynomial function, and develop an individual tree mortality model based on a logit function.

**Material and methods:** The study took place in a mixed beech-hornbeam naturally regenerated stand in Hesse forest (NE of France), where a square design 42*42m comprising 64 square plots of 5.25m side was inventoried each year since 2001 until 2012. Measurements included tree species and status (dead or alive), girth at breast height for all trees and total height for a sample of living trees (beech and hornbeam). The size-density trajectories of the 64 plots describing the course of the number of living trees in relation with the mean stand girth, in logarithmic scales, were modeled with a piecewise polynomial function fitted with a mixed-effects model. On the other hand, the individual tree mortality model was fitted with a logit function including several independent variables defined at tree, plot and stand levels.

**Results:** The size-density trajectory of mixed beech-hornbeam naturally regenerated stands was successfully fitted using the same function as for pure stands, with a plot-level random component that appeared linearly related to site fertility, initial density of trees (N_0_) and relative initial proportion of beech trees. The mortality onset appeared to occur at a higher density (RDI = 0.5) in mixed beech-hornbeam naturally regenerated stands than in pure beech even-aged stands (RDI = 0.29), while the maximum density (RDI = 1) was reached at a comparable relative number of surviving trees (N/N_0_). An individual tree mortality model could also be fitted using a logit function. The probability of mortality of trees appeared linked to individual (social status, species), collective (relative density) and site (fertility, water stress) factors.

**Conclusion:** The size-density trajectory model first developed for pure even-aged stands appeared well adapted to mixed beech and hornbeam natural regenerations, and the individual tree mortality model constructed at the same time for these stands revealed that water stress induced mortality was relatively comparable for beech and hornbeam. These two models give the opportunity to simulate the development of mixed beech and hornbeam regenerations with the addition of a growth model.

## 1 Introduction

For naturally regenerated stands, the different stages in the development following the regeneration phase are crucial as they greatly influence the future of the stand, in terms of species mixture, productivity and wood quality. In this scope, size-density management diagrams have been developed – more often for pure stands – to define stand density dynamics, especially in the US and Canada (Penner et al. 2006; Vanderschaaf et al. 2012) and more rarely in Europe (Barrio Anta and Alvarez Gonzalez 2005, Lopez-Sanchez and Rodriguez-Soalleiro 2009; Patricio et al. 2017). These are based on size-density relationships that allow the prediction of mortality during the whole stand development, and then help the forest manager to anticipate mortality through cleaning or thinning operations (Barrio et al. 2005).

Size-density trajectory models have been developed abroad for many years (Puettmann et al. 1993) and in France in recent years for several species growing in pure even-aged stands (Ningre et al. 2016a, 2016b; Ningre et al. 2019; Le Goff et al. 2021). The remarkable feature of these trajectories, in which the dynamics of stands of different initial densities are plotted on double logarithmic scales (Ln(Cg), Ln(N)), where Cg and N are respectively the mean girth and the total number of trees of the stand, is that a trajectory is entirely determined by the initial density (N_0_) of the stand.

Beech-based natural regenerations, often mixed with other species, in particular hornbeam (Morneau et al. 2008; Ratcliffe et al. 2015; Dietz et al. 2021), but also oak and other species like pioneer species, would need the development of this kind of silvicultural tools. How these mixtures will develop and how will they become sustainable stands, in particular dealing with climate change ? As a matter of fact, beech and hornbeam are more sensitive to drought than oak (Bréda et al. 2006; Friedrichs et al. 2009), while hornbeam is also considered to tolerate single years of severe drought (Leuzinger et al. 2005). So, the future of these beech-based mixtures remains uncertain in the context of more intense and/or more frequent spring and summer droughts (Jump et al. 2006; Geßler et al. 2007).

The construction of size-density relationships for these mixed beech-based natural regenerations should be adapted from similar recent studies dealing with pure stands, by taking account, if necessary, of the mixture of species. Moreover, an individual mortality model could allow to predict which trees will die, in terms of tree dimensions and species (Maleki et al. 2015; Wunder et al. 2007), and then to inquire about stand structure.

To this end, we took advantage of an instrumented experiment belonging to the European flux-monitoring network (CarboEurope IP) and set up in 2001 in a mixed beech natural regeneration in Hesse forest (NE France) to install experimental plots with annual tree monitoring. In this lowland beech forest with varying local density and species composition (Longdoz et al. 2003), mortality and growth were recorded on these plots over a twelve years period. Moreover, we controlled environmental factors, such as climate and soil characteristics (Quentin et al. 2001) that may be involved in individual tree mortality (Crouchet et al. 2019; Qiu et al. 2015; Ruiz-Benito et al. 2013) and in the course of stand size-density trajectories.

The two main objectives of this study were then to develop size-density relationships for mixed stands of beech and hornbeam, the two principal species of our experimental plots of Hesse forest, and to establish an individual tree mortality model specific of each species in relation with its requirements in terms of density of trees, species diversity and environmental conditions. The possibility of using the individual tree mortality model to predict the development of mixed beech-hornbeam stands will also be mentioned, as it can appear for some authors as a constraint in growth models together with the use of size-density relationships (Monserud et al. 2005; Yang and Titus 2002).

## 2 Material and Methods

### 2.1 Experimental design

The study was carried out in the state forest of Hesse in the North-East of France (48°40′27′′ N; 7°03′53′′ E, altitude 305 m). It is a high forest, naturally regenerated, and composed mainly of oak (*Quercus petraea* and *Q. robur*, 40%) and European beech (*Fagus silvatica* L., 37%), the remaining species being mostly hardwoods (23%). The climate is continental with oceanic influences: the mean annual temperature averages 9.2°C and total annual precipitation 820 mm.

The experimental instrumented site, named “Hesse-2”, was installed in 2001 near the middle of a 15-ha naturally regenerated mixed broadleaved stand, twelve years old, and was surrounded by a fence installed at 1.6 m inside the site limits. The soil, moderately deep is classified as *luvisol* covered by a humus of type *oligo-mull* (Ngao et al. 2007). This site was primarily devoted to ecophysiological studies, notably the study of the carbon balance of the stand, but also the study of the “differential response to soil drought among co-occuring broadleaved species” (Zapater et al. 2013).

The study of the stand density dynamics was conducted in the 64 plots (squares of 5.25 m size) comprising the experimental site (Fig. 1). The dimensions of the design (42*42m) exceeded that of the fenced area (40.4*40.4m) and was installed at the side of a silvicultural trail on two opposite sides of the experimental design.

**Fig. 1.**
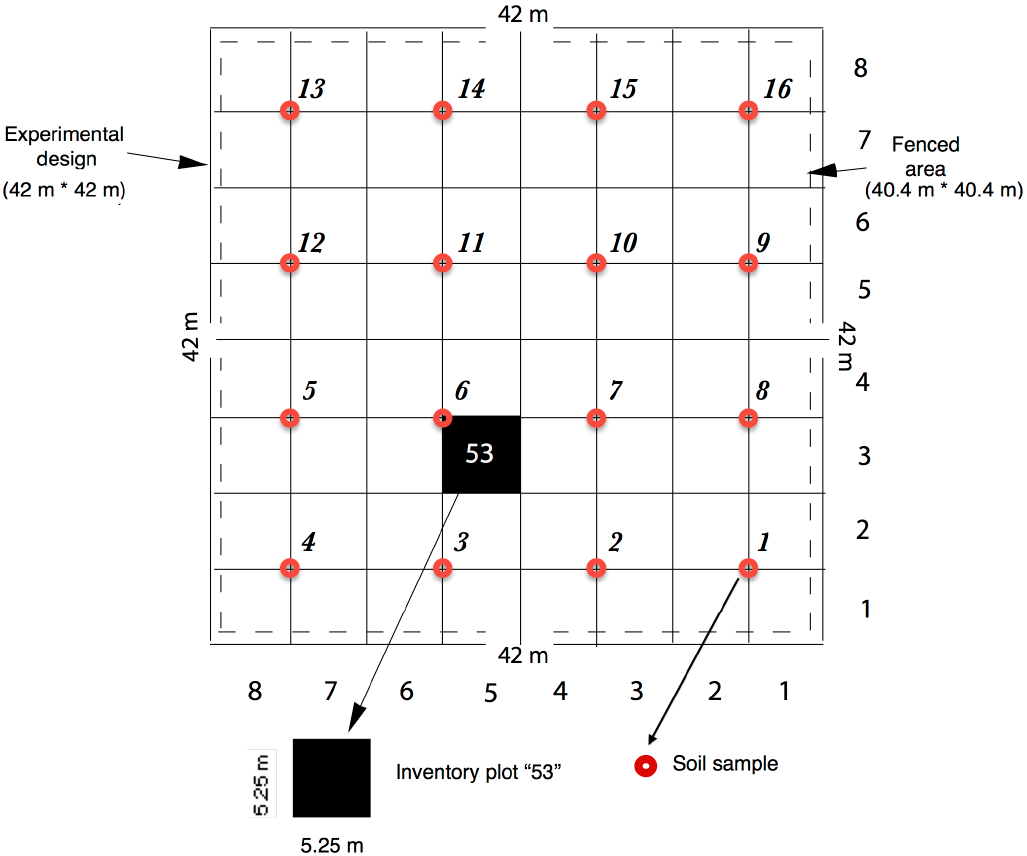
Experimental design: square area of 0.1764 ha comprising 64 square plots with sides measuring 5.25 m and identified with a number composed of column number (1 to 8) followed by line number (1 to 8): as an example, the plot 53 is situated on the map. Red circles, numbered 1 to 16 on the map are the locations where soil samples were planned

### 2.2 Tree measurements

Tree inventory was conducted annually in each of the 64 “inventory plots” from year 2001 until year 2012^1^. The following characteristic variables were recorded for each tree: tree species, tree status (dead or alive), tree diameter (d130) or girth (c130) at breast height (cm). In addition, each year, tree height (ht, m) was measured with a telescopic pole for a representative sample of the three major tree species present in the experimental stand. The number of sampled trees depended on the relative frequency of each species in the stand: beech (*Fagus silvatica* L., 50 trees sample), hornbeam (*Carpinus betulus* L., 30 trees sample) and oak (*Quercus petraea*, 19 trees sample, from year 2005 only). Height-girth curves were fitted^2^ for each species to (c130, ht) sample data allowing calculating the mean height (Hg) of the stand component of mean girth Cg (Table 1). Finally, all trees were numbered from year 2005, allowing following annual tree girth increment and individual tree mortality for the period 2005-2012.

**Table 1.** Stand characteristics of living trees in 2001 for the three major species and for all species together in the experimental site *Hesse-2*: number of trees per ha (*N/ha*) and basal area per ha (*G/ha*), mean girth (*Cg*) and height (*Hg*) (dead trees unregistered in 2001)

| Species | N/ha | N/ha (%) | G/ha | G/ha (%) | $C_g$ (cm) | $H_g$ (m) |
| --- | --- | --- | --- | --- | --- | --- |
| Beech (1) | 7035 | 41.9 | 6.81 | 66.0 | 11.03 | 4.95 |
| Hornbeam (2) | 8124 | 48.4 | 2.45 | 23.7 | 6.15 | 4.01 |
| Oak (3) | 794 | 4.7 | 0.63 | 6.1 | 9.95 | - |
| (1)+(2)+(3) | 15953 | 95.1 | 9.89 | 95.8 | 8.83 | - |
| All species | 16774 | 100 | 10.32 | 100 | 8.79 | - |

### 2.3 Stand characteristics

Tree inventories allowed to calculate the principal stand characteristics at installation of the experimental stand in 2001 and for successive years until 2012, for the 64 experimental plots and for the whole stand considered as the grouping together of the 64 plots: number (N) and basal area (G) per hectare of live and dead trees for each species present, mean girth (Cg) and height (Hg) for the three main species represented (beech, hornbeam and oak). These species totalize 95% and 96 % of the total number of trees and total basal area respectively (Table 1). At the first survey in 2001, a total of about 3000 trees were measured. Among the three main species, beech and hornbeam were, by far, the most important components of the stand. Beech trees were slightly fewer but had a larger average size Cg than hornbeam trees, and then contributed more to stand basal area. Then, at the installation of the experiment (2001), beech appeared as the dominant species in the stand in terms of size and height, and hornbeam in terms of frequency. The other species present in the stand were the following: silver birch (*Betula pendula*), wild cherry (*Prunus avium*), aspen (*Populus tremula*), and goat willow (*Salix capreae*).

A diversity index was calculated to represent the diversity of species at stand and plot levels: we used the *Simpson index* (Ind.div) defined as 1/∑p_i_^2^ where p_i_ is the proportion n_i_/N of the trees of each species *i* represented (Cordonnier et al. 2012). Three “species” were considered to calculate the Simpson index: “beech”, “hornbeam” and “other species” considering the preeminence of beech and hornbeam in the stand (Table 1). Then, the diversity index is equal to 1 (minimum value) if there is only one species present in the plot, and it peaks to 3 (maximum value) when the three main “species” are equally represented.

Moreover, a stand density index (*RDI*) was calculated to characterize the relative density of each of the 64 plots^3^. If *N*_*i*_ is the number of trees per ha of a given plot *i* and *Cg*_*i*_ its mean girth, then:

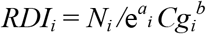

where *a*_*i*_ is the intercept and *b* the slope of the maximum size-density line of plot *i*, Ln(*N*_*i*_) = *a*_*i*_ + *b* Ln (*Cg*_*i*_), with *a*_*i*_ = *a* + *∂*_*i*_; *a* is the mean intercept parameter of this line and *∂*_*i*_ the random effect on *a* due to plot *i*; *b* is the fixed parameter of the maximum size-density lines, common to all 64 plots^3^. The choice of *b* as a fixed parameter and of *a* as a random parameter were suggested by the observation of the size-density trajectories of the 64 plots (see *Results* section).

The large variability of stand characteristics at plot level is shown in the following table (Table 2).

**Table 2.** Variability of the main stand characteristics (N, G, Cg, RDI, Ind.div) among the 64 plots of the experimental site “Hesse-2” in 2001

| Characteristic | Minimum | Mean | Maximum |
| --- | --- | --- | --- |
| N ($\text{ha}^{-1}$) | 2902 | 16774 | 66757 |
| G ($\text{m}^2 \text{ha}^{-1}$) | 4.34 | 10.32 | 20.85 |
| Cg (cm) | 4.69 | 9.64 | 17.40 |
| RDI | 0.22 | 0.46 | 0.84 |
| Ind.div | 1.00 | 1.80 | 2.93 |

### 2.4 Environmental measurements

#### 2.4.1 Climate

Climatic data were obtained from the site “Hesse-1”, a nearby and older experimental site in Hesse forest, at the scale of the day, which allowed the calculation of precipitation (*P*, mm) and global radiation at month level during the studied years (Zapater 2009, 2013). This, in turn, allowed the calculation, at year level, of the potential evapotranspiration (*ETP*, mm) and of the real evapotranspiration (*ETR*, mm) for a defined soil water holding capacity (*RU*) fixed hypothetically to 150 mm. The variations of these major climatic parameters for the studied period in Hesse forest appear in Fig. 2A: in particular, 2003 was by far the driest year in the studied period, leading to a reduced ETR in spite of a high ETP that year.

**Fig. 2A.**
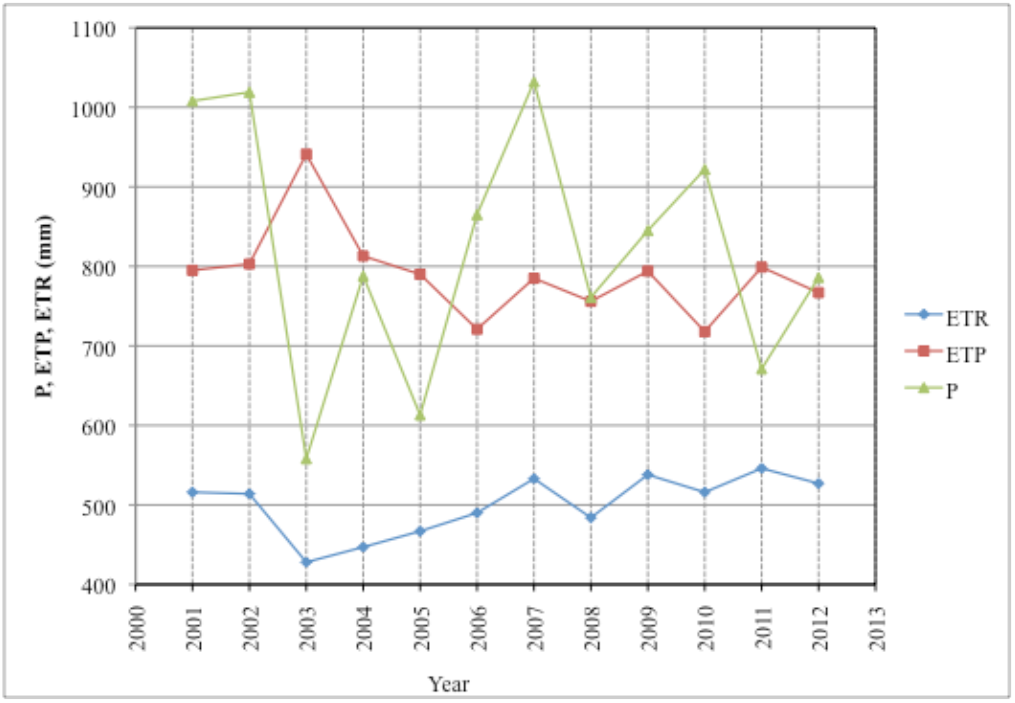
Evolution of the principal yearly climatic variables considered (ETR, ETP, P) for the site ‘Hesse-2’ in the studied period.

**Fig. 2B.**
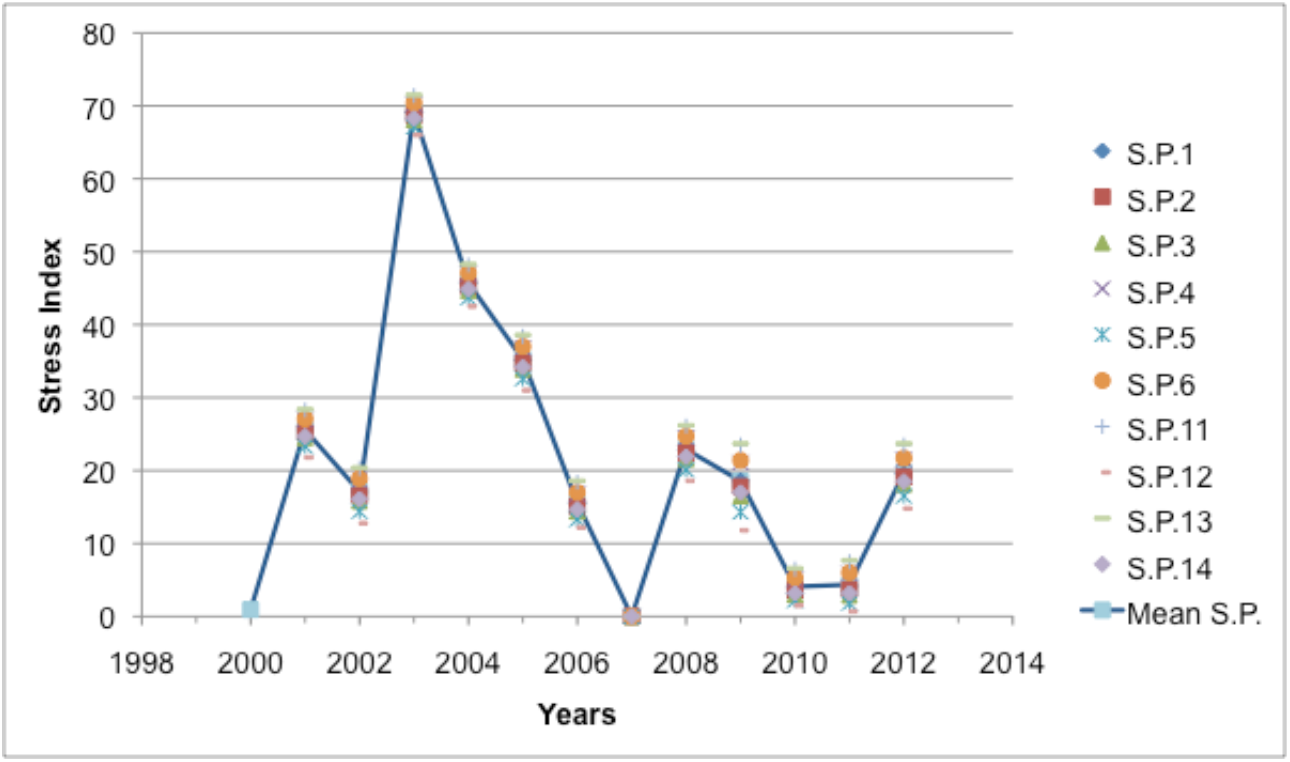
Evolution of the yearly mean *water stress index* (Mean S.P.) for the period 2000 to 2012: the yearly values of the stress indexes of the 10 sample points retained (S.P.), also shown on the graph, were calculated with the model “*Biljou*”

#### 2.4.2 Soil

Soil conditions were described for clusters of 4 inventory plots (see Fig. 1) after extracting a core sample of soil at the center of each cluster called “sample point”. The texture and thickness of the different soil layers were recorded from the core samples in order to estimate the water holding capacity of the soil (RU, mm) at each sampling point. Soil texture, appreciated with a sensitive touch, allowed determining the texture of each layer using the *soil texture triangle* based on percentages of sand, loam and clay (Duchaufour, 1997). The thickness of each layer was determined after taking account of the soil compactness due to an increase of clay content and of the percentage of stones that both limit root development: these two limits to root development defined the potential depth of soil useful for fine root development. The depth at which occurs the *BT horizon* (see Baize and Girard, 2008), characterized by a sharp increase of the clay content, was also recorded: it coincides with a drastic reduction of root density (see root profiles in Zapater, 2009, page 38).

Then, the water holding capacity (in mm) of each layer was estimated from its thickness and potential water capacity (in mm/cm) determined by its texture (data from Baize D. in Ridremont et al. 2012, p25). Adding the water capacity of each soil layer allowed to calculate the water capacity of the entire soil profile at each considered sample point^4^ (see Fig. 1). The characteristics of the soil limiting root development (depth of *BT* layer) and determining water availability (*RU*) for trees at each sample point considered appear in Table 3. So, the depth where began the *BT* layer was comprised between 24 and 54 cm and varied by twice as much, whereas the estimated RU at sample points was comprised between 115 and 145 mm (Table 3).

**Table 3.** Depth of soil *BT* layer limiting root development and water holding capacity (*RU*) at each sample point considered in the study

| Soil characteristics | Sample points (see Fig. 1) |  |  |  |  |  |  |  |  |  |  |
| --- | --- | --- | --- | --- | --- | --- | --- | --- | --- | --- | --- |
|  | 1 | 2 | 3 | 4 | 5 | 6 | 11 | 12 | 13 | 14 | Mean |
| *BT* layer (cm) | 24 | 29 | 41 | 34 | 44 | 54 | 43 | 40 | 33 | 37 | 39.4 |
| *RU* (mm) | 129.8 | 132.9 | 135.3 | 123.2 | 145.1 | 122.1 | 115.9 | 147.5 | 115.3 | 134.1 | 130.2 |

#### 2.4.3 Water stress index

A *water stress index*^*5*^ could then be calculated for each sample point and for each year considered^6^ using “*Biljou”*, a forest water balance model (Granier et al. 1999, 2011; *Biljou* 2014) (Fig. 2B). As some sample points were missing, the yearly mean values of stress indexes were retained for the 64 experimental plots, considering the small variation of the stress index^7^ with the water holding capacity of the soil^8^ in our conditions (Fig. 2B).

### 2.5 Methods

#### 2.5.1 Stand dynamics

Stand dynamics (growth and mortality) was studied from year 2001 to year 2012 by using a size-density trajectory model expressing the logarithm of the number of trees N as a function of the logarithm of the mean girth Cg. The size-density modeling framework developed for pure even-aged stands of beech (Ningre et al. 2016a), oak (Ningre et al. 2019), Douglas-fir (Ningre et al. 2016b) and ash (Le Goff et al. 2021) was applied in this study to represent the size-density trajectories for the mixed stands in the 64 experimental plots. As for ash, due to the scarcity of data (no replication for each plot), a global fit of the 64 size-density trajectories was performed.

The evolution of the number of trees per hectare (N), before mortality onset until maximum density has been reached and next, was modeled by the following family of continuous smoothed functions *f* of *Cg*, depending only on the four parameters *Cg*_*0*_, *N*_*0*_, *a* and *b*:

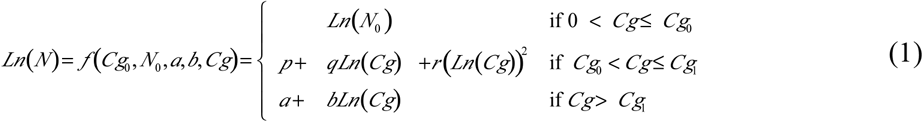

Where:

- *Cg*_*0*_ is the mean girth at breast height of a stand at the onset of regular mortality, *N*_*0*_ its number of trees per hectare prior to this onset^9^ and *Cg*_*1*_ is the stand mean girth at breast height when the maximum RDI is reached^10^
- *a* and *b* are the parameters of the maximum size-density line :

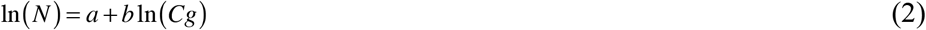

*p, q* and *r* are given by the following equations (Le Goff et al. 2021):

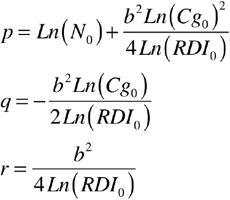

with *RDI*_*0*_, the density at mortality onset, equal to 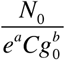

It follows that Equation (1) only depends on the parameters *Cg*_*0*_, *N*_*0*_, *a*, and *b* because of the continuity constraints on this equation and its derivative (Ningre et al. 2016, 2019; Le Goff et al. 2021).

We could satisfactorily fit Equation (1) globally to the trajectory data of the 64 plots, adding to the model the constraint of inflection points aligned^11^ parallel to the maximum size-density line as was observed for beech, oak and ash previously (Ningre et al. 2019, Le Goff et al. 2021), that is:

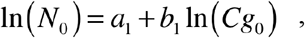

with *b*_*1*_ = *b* and *a*_*1*_ = ß *a* (ß is a coefficient of proportionality)

Equation (3) was fitted using a mixed-effects model where *b* and ß were fixed parameters with *b* common to all 64 plots and *a*_*1*_ had a mean value *a* common to all plots with an added normal random effect (∂_i_) due to variability among the 64 plots. The choice of *b*_*1*_ as a fixed parameter equal to *b* and of *a*_*1*_ as a random parameter were suggested by the observation of the size-density trajectories of the 64 plots (see *Results* section). In the model, *N*_*0*_ was introduced as an independent variable. The resulting statistical equation was the following:

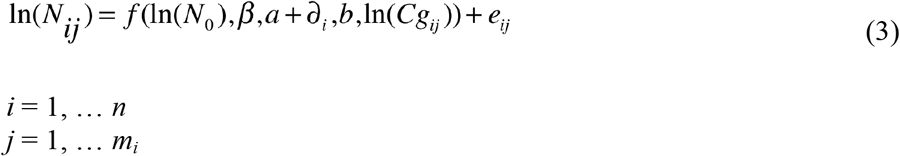

In Eq. 3, the indices *i* and *j* stand respectively for plot (1 to 64) and year (2001 to 2012, for a large majority of the 64 plots), n is the number of plots (n=64), *m*_*i*_ is the number of years when plot *i* was measured. The residual errors *e*_ij_ are independent, normally distributed random variables with mean 0.

If *N*_*i*_ is the number of trees per ha of a given plot i and *Cg*_*i*_ its mean girth, then:

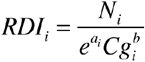

where a_i_ = a + ∂_i_

In addition, relations between parameters of the size-density trajectory model and environmental factors characterizing each plot (Table 4) were investigated.

**Table 4.** List of explanatory variables (at tree, plot and stand levels) considered in the logit regression model explaining tree mortality in the period 2005-2012. The status (*st*)^12^ of the tree is the dependent variable to be explained.

| Variable | Explanation | Min | Max | Unit |
| --- | --- | --- | --- | --- |
| **Tree level** |  |  |  |  |
| *Tree species (S)* | Species considered: beech ($S_b$), hornbeam ($S_h$), “others ($S_o$)” (= silver birch, wild cherry, aspen, goat willow, oak) | S = 1 if the tree is of the concerned species, 0 if not | | - |
| *c130* | Girth at breast height | 0.0 | 77.1 | cm |
| *c130/Cg* | Ratio of tree girth to mean girth (social status) | 0.0 | 4.72 | - |
| **Plot level** |  |  |  |  |
| *N* | Number of living trees | 1451 | 60227 | Nb ha<sup>-1</sup> |
| *N<sub>0</sub>* | Initial<sup>13</sup> number of trees | 2902 | 67846 | Nb ha<sup>-1</sup> |
| *N/N<sub>0</sub>* | Relative number of living trees | 0.18 | 1.04 | - |
| *Nh<sub>0</sub>/N<sub>0</sub>* | Initial proportion of alive beech trees | 0.0 | 100.0 | % |
| *G* | Basal area of living trees | 4.34 | 44.72 | m<sup>2</sup> ha<sup>-1</sup> |
| *G<sub>0</sub>* | Initial basal area of living trees | 4.34 | 20.85 | m<sup>2</sup> ha<sup>-1</sup> |
| *Cg* | Mean girth at breast height | 4.69 | 39.26 | cm |
| *C0* | Dominant girth at breast height in 2010 | 29.6 | 72.2 | cm |
| *C0(2012)* | Dominant girth at breast height in 2012 | 31.0 | 77.1 | cm |
| *H0* | Dominant height in 2010 | 11.3 | 15.3 | m |
| *RDI* | Relative stand density | 0.21 | 1.66 | - |
| *Ind.div<sub>0</sub>* | Initial diversity index | 1.0 | 3.0 | - |
| *BT* | Depth of appearance of the soil layer *BT* | 24.0 | 54.0 | cm |
| *RU* | Water holding capacity of the soil | 115.3 | 147.5 | mm |
| **Stand level** |  |  |  |  |
| *Stress<sub>j</sub>* | Water stress index for year *j* (same for each plot as water stress slightly dependent of *RU* and *RU* varies only slightly among the plots) | 0 | 63.6 | - |
| *Stress<sub>j-1</sub>* | Water stress index for year *j-1* (same for each plot, as for year *j*) | 0 | 63.6 | - |

#### 2.5.2 Individual tree mortality

The probability of mortality (*Pm*) of a given tree can be defined by the following function *P*:

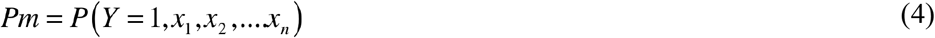

where Y is the status of the tree (1 if dead; 0 if alive) and X(x_1_, x_2_, … x_n_) is the matrix of predictor variables x_i_.

The logit function (Maleki & Kiviste, 2015) was used to explain the probability of mortality *Pm*, that is:

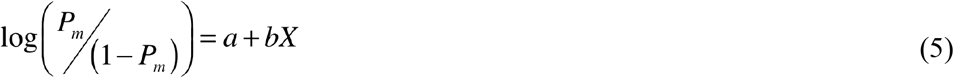

that corresponds to:

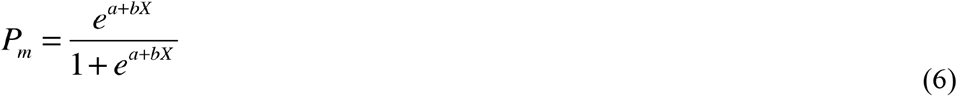

For each tree and each year considered (from 2005 to 2012) potential explanatory variables of the annual probability of mortality were selected and calculated at tree, plot and stand levels (stand considered as the grouping together of the 64 plots). The values of the variables introduced into the model are those for which tree status is observed, except for C0 that was calculated for two reference years (2010 and 2012) and for H0 calculated only for one year (2010). The water stress index of the year preceding the mortality survey (stressj-1) was also considered (Table 4). In fact, the climatic conditions of the previous autumn, especially water balance, influence current early wood growth of several species (Lebourgeois et al. 2005) and can consequently be a factor of tree mortality.

Equation (5) was explored using *DataDesk* linear regression software (*DataDesk 6*.*3, 2011*) with potential explanatory factors introduced successively as independent variables. The residuals of each fitted equation were plotted against the remaining factors selected to detect any other dependence of tree mortality. Finally, Equation (5) was fitted using a logistic regression model implemented in “*R”* (R Development Core Team, 2012), with “*ROCR*” package for calculating the quality criteria of the fitting (Wunder et al. 2007).

### 2 Results

#### 3.1 Size-density trajectories

Equation (3) representing the size-density trajectories of plots was fitted with *N*_*0*_ as an independent variable and assuming that there was a random effect on parameter *a* due to variations between plots. In addition, the correlation between yearly data plots was taken into account in the residual errors (correlation structure “ARMA” in R (2012), *nlme*). Table 5 gives the parameters and statistics of the fit of Eq. (3), and the mean size-density trajectory, with parameters *a, b* and *ß* (with *a*_*1*_ = *ß a*), is represented in Fig. 3, together with the observed size-density data of the 64 plots^6^.

**Table 5.** Parameter estimates and statistics of Equation (3) fit to the mixed beech-based stand data of the 64 plots of Hesse-2

| Parameter | Estimate | Std. Error | *t* value | *P* value |
| --- | --- | --- | --- | --- |
| *a* | 14.342 | 0.207 | 69.10 | $\leq 0.0001$ |
| *b* | -1.783 | 0.066 | -26.94 | $\leq 0.0001$ |
| $\beta$ | 0.952 | 0.004 | 218.12 | $\leq 0.0001$ |
| Residual Std. Deviation = 0.087, with n = 624 degrees of freedom |  |  |  |  |

**Fig. 3.**
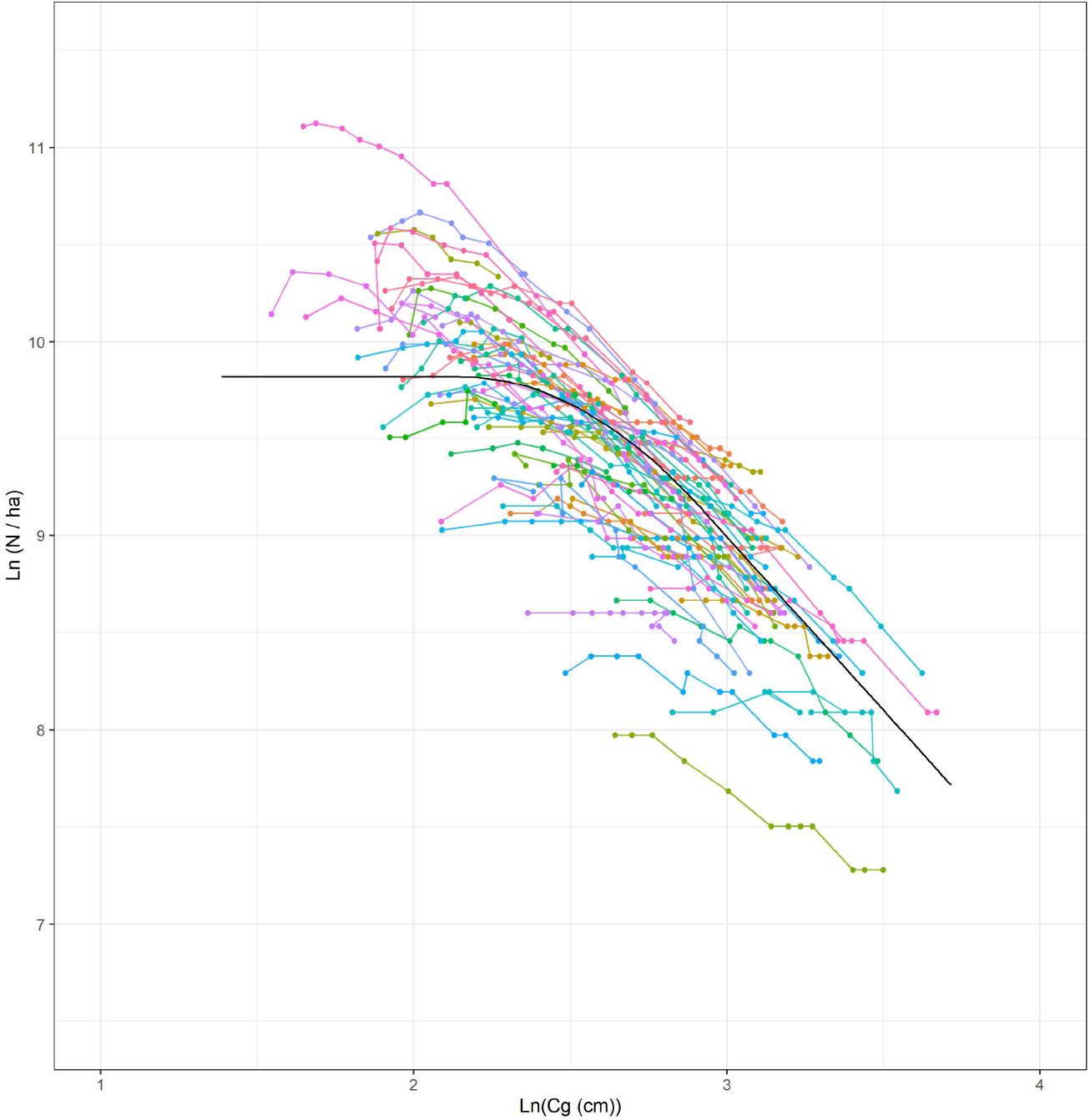
Observed size-density trajectories of the 64 plots of the experimental site “Hesse-2” (colored lines), together with the mean fitted trajectory (black line) based on Eq. (3)

Size-density trajectories resulting from the fit of Eq. 3 are shown separately for each plot and compare favorably with the observed plot data (Fig. 4). The normalized residuals, corrected to allow for intra-plot auto-correlation, did not show time dependence (Annex, Fig. 5) and appeared unbiased with no marked heteroscedasticity (Annex, Fig. 6). Moreover, the normality of the within-plot errors appeared satisfied (Annex, Fig. 7) as that of the random effects applied on parameter *a* (Annex, Fig. 8).

**Fig. 4.**
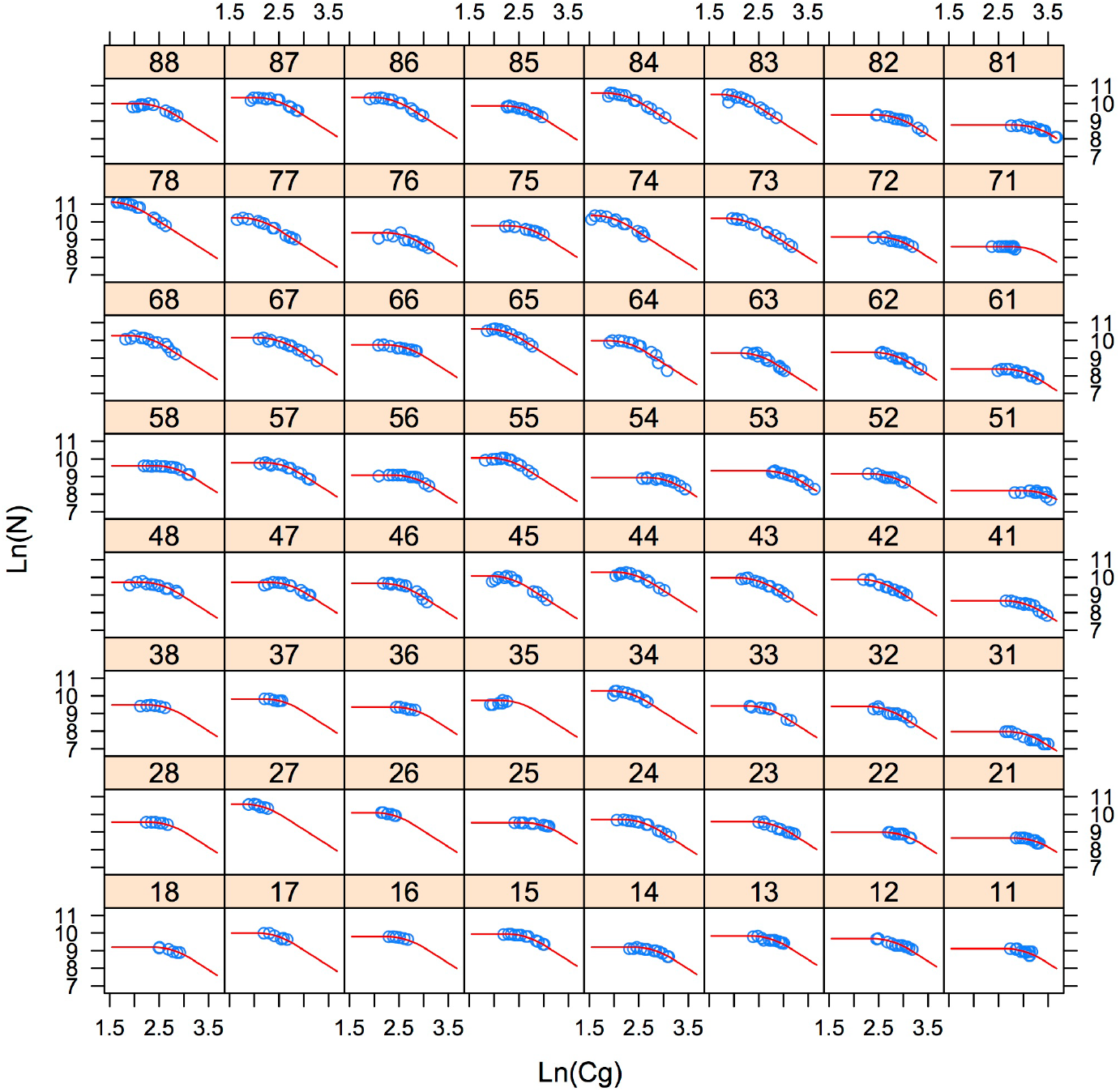
Fitted size-density trajectories of the 64 plots of the experimental stand of Hesse-2 based on Eq. (3) and arranged as in the field, together with the observed size-density data of each plot

**Fig. 5.**
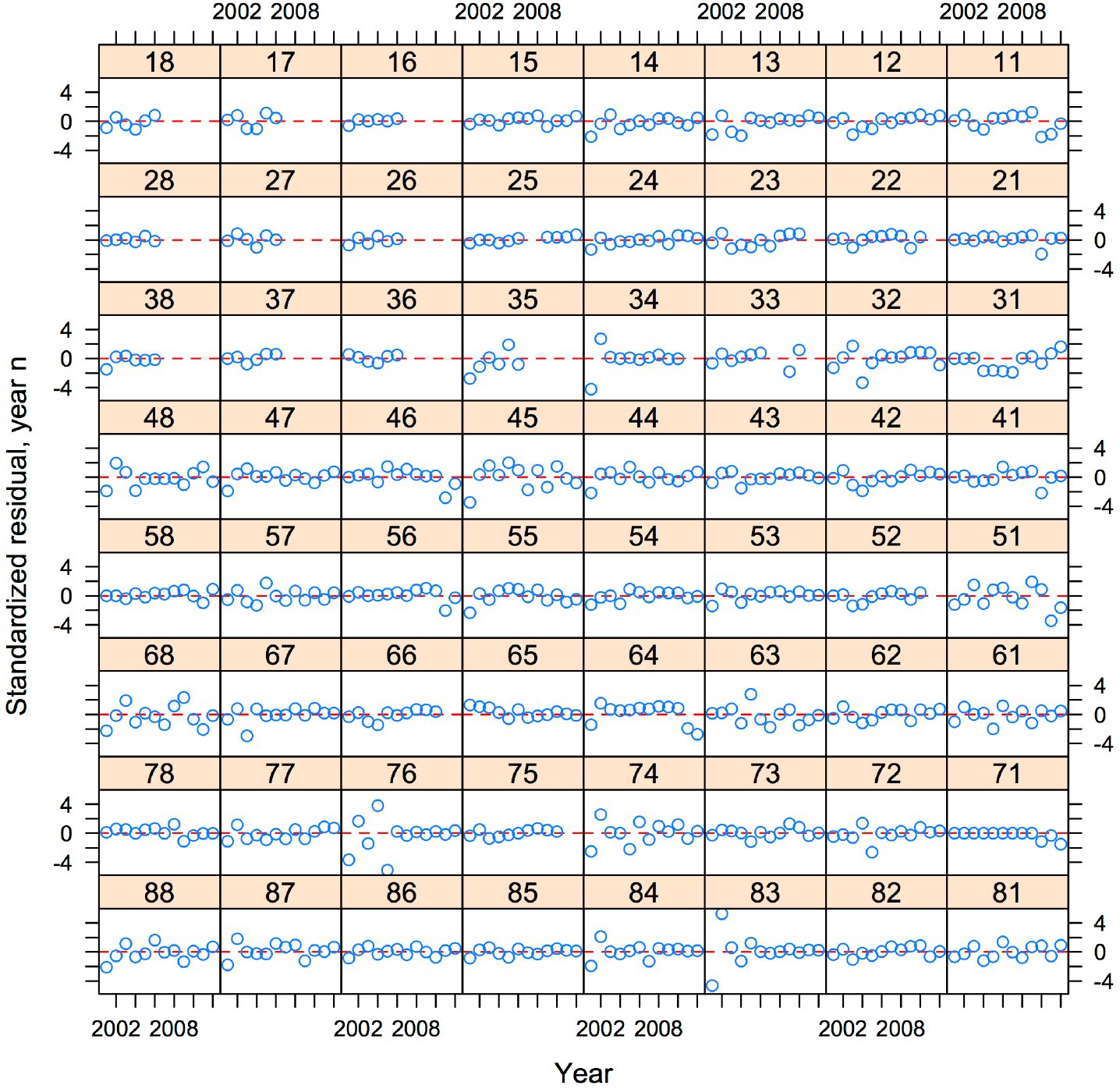
Auto-correlation plot showing that the one-year lag plots of the normalized residuals did not show evidence of year-to-year correlation patterns of the within-plot errors

**Fig. 6.**
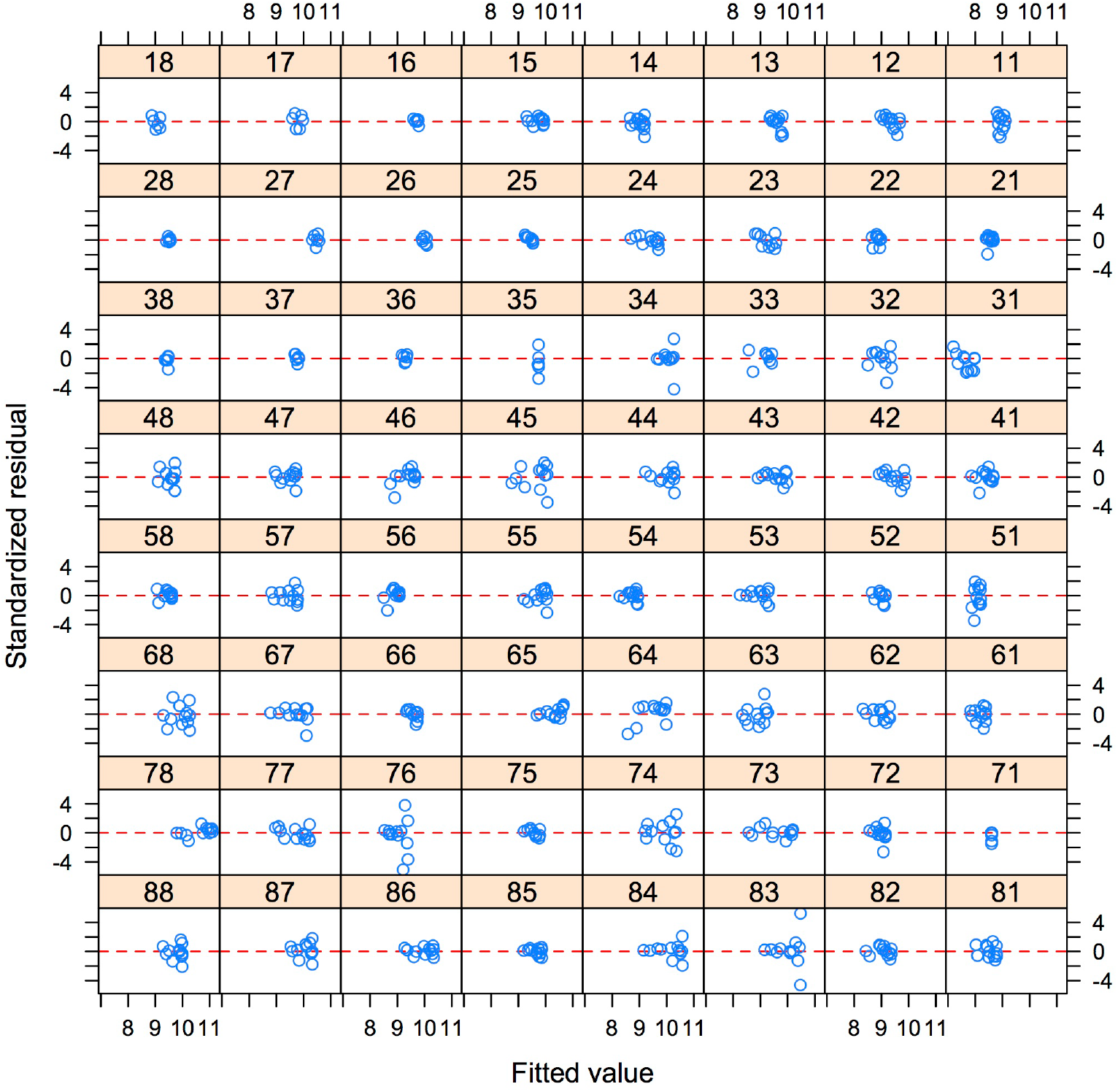
Scatter plot of the standardized residuals versus the fitted values from the size-density fit of Equation (3) to the data of the 64 plots of Hesse-2 experimental stand

**Fig. 7.**
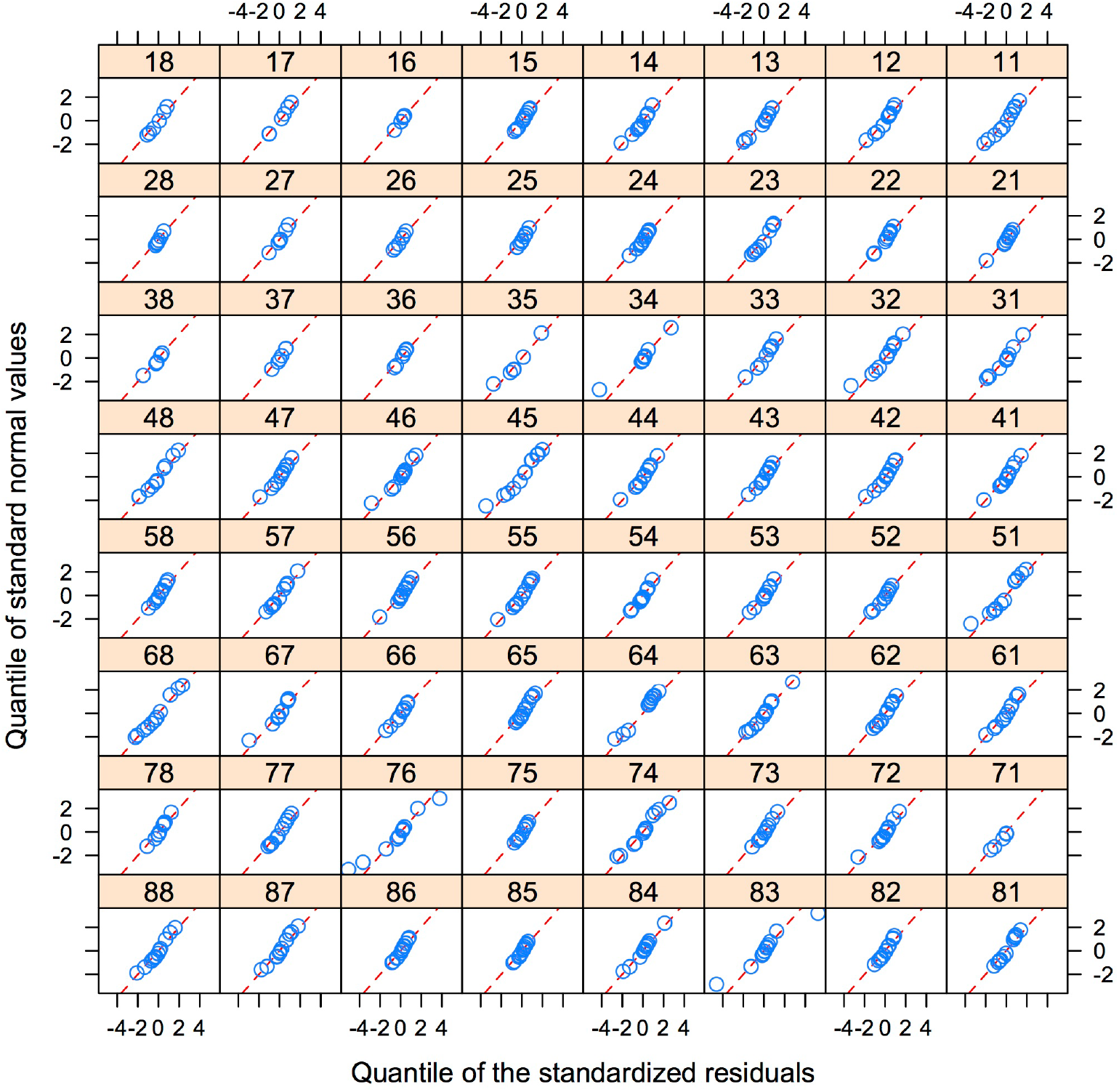
Normal (QQ) plot of the normalized residuals of the fit of Eq. 3 to the 64 plots of the beech-based experimental stand of Hesse-2. The linearity of the relationships provides evidence for the normality of the within-plot errors

**Fig. 8.**
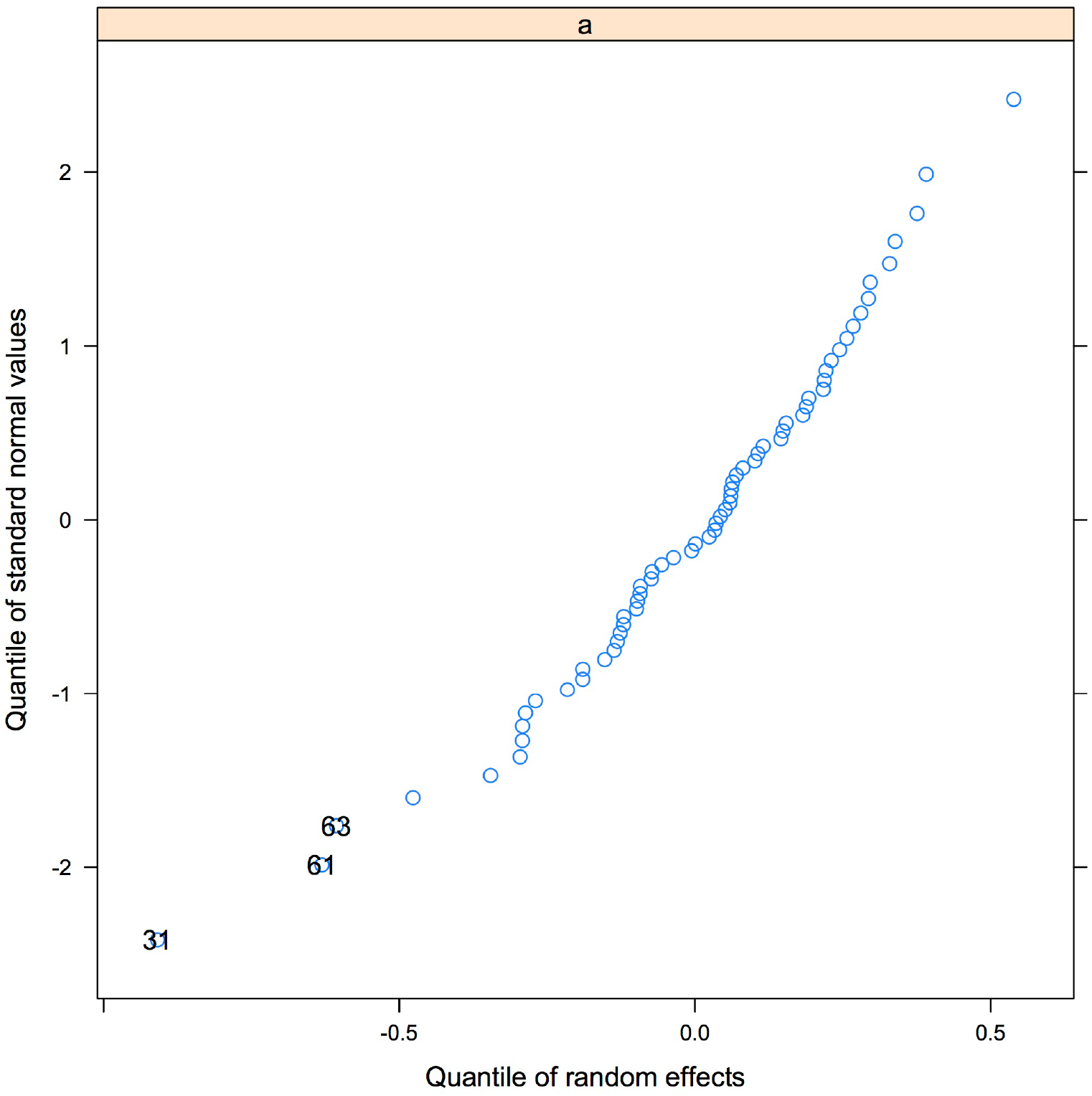
Normal (QQ) plots for the random effects on parameter *a* assessing their normality: three points only – corresponding to plots 31, 61 and 63 – show a minor deviation from the straight line depicted by the other points

Stand characteristics of the size-density trajectories at the critical stand development stages corresponding to the onset of mortality and reach of maximum RDI, respectively (Cg_0_, RDI_0_) and (Cg_1_, N_1_), were derived from Equation (3) for the 64 plots as a whole (Table 6).

**Table 6.** Estimations of the size-density trajectory characteristics from N_0_ and Eqs. (3), (7) and (8), at mortality onset (Cg_0_, RDI_0_) and reach of maximum density (Cg_1_, N_1_), for the mean of the 64 plots of Hesse-2

| Parameter | Mortality onset |  |  | Reach of maximum RDI |  |
| --- | --- | --- | --- | --- | --- |
| | $N_0^{(1)}$ (ha<sup>-1</sup>) | $Cg_0$ (cm) | $RDI_0$ | $Cg_1$ (cm) | $N_1$ (ha<sup>-1</sup>) |
| Estimate | 18413 | 8.57 | 0.50 | 18.55 | 9251 |

<sup>(1)</sup> $N_0$ is the mean of the initial number of trees per ha observed in the 64 subplots and introduced in Eq. (3) as an independent variable

RDI_0_, the relative density at mortality onset, is equal to N_0_/e^a^Cg_0_^b^. Cg_1_, the mean girth when the maximum RDI is reached, is obtained by the following equation (Ningre et al. 2019; Le Goff et al. 2021):

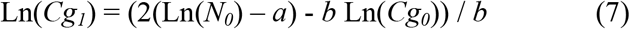

*N*_*1*_ is then obtained from the maximum size-density equation, that is:

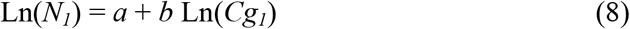

### 3.2 Effects of environmental variables on size-density trajectories

The possible effects of environmental variables on the size-density trajectories were analyzed through relationships (if any) between the random parameter ∂_i_ of Eq. 3, and retained variables at plot *i* level, that is fertility indexes calculated for year 2010 (*H0*) and years 2010 and 2012 (*C0*), water holding capacity of the soil (*RU*), depth (*BT*) of the BT layer, and initial stand characteristics before mortality appears (basal area (*G*_*0*_), number of trees per ha (*N*_*0*_), mean girth (*Cg*_*0*_), diversity index (*Ind*.*div*_*0*_), relative basal area (*G/G*_*0*_) and number of beech trees per ha (*Nh*_*0*_*/N*_*0*_)).

The random parameter ∂_i_ appeared linearly related to Ln(*N*_*0,i*_) and Ln(*Cg*_*0,i*_) through the following equation:

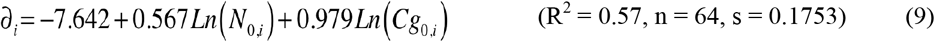

The dependences of ∂_i_ on Ln(*N*_*0,i*_) and Ln(*Cg*_*0,i*_) are illustrated by Fig. 9 where partial residuals of Eq. 9 were plotted. Moreover, the following equation (Eq. 10)^7^ could be established:

**Fig. 9.**
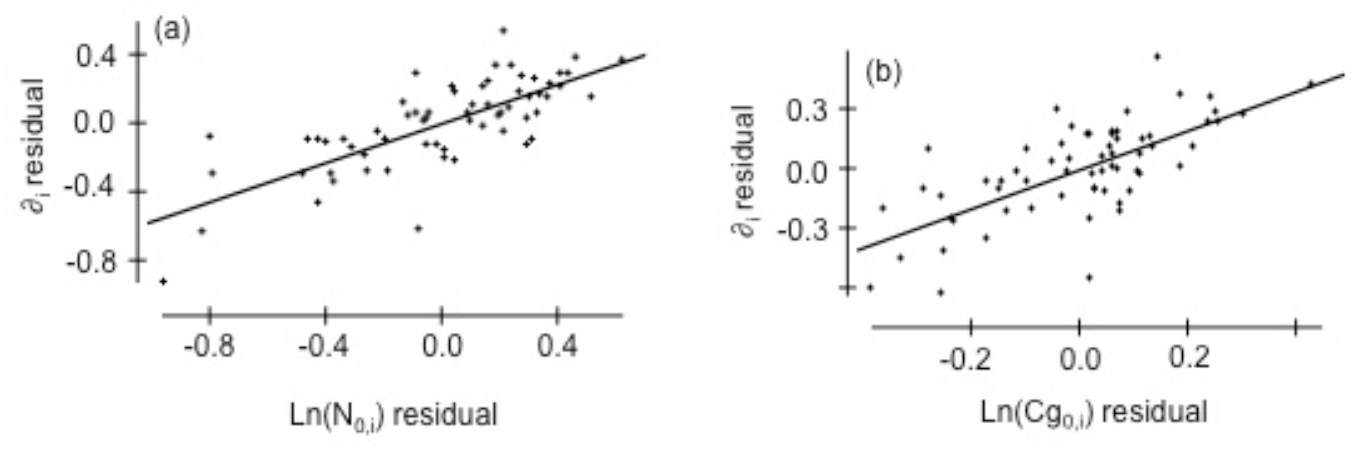
Linear dependences between the random effect *∂*_*i*_ on parameter *a* in Equation (3) and two stand variables: (a) initial tree number (*N*_*0,i*_) and (b) mean initial girth (*Cg*_*0,i*_), variables which are log-transformed; the two graphs show the linear dependences between partial residuals of ∂_i_ versus each of the two log-transformed variables (*N*_*0*_ and *Cg*_*0*_) in Eq. (3)

**Fig. 10.**
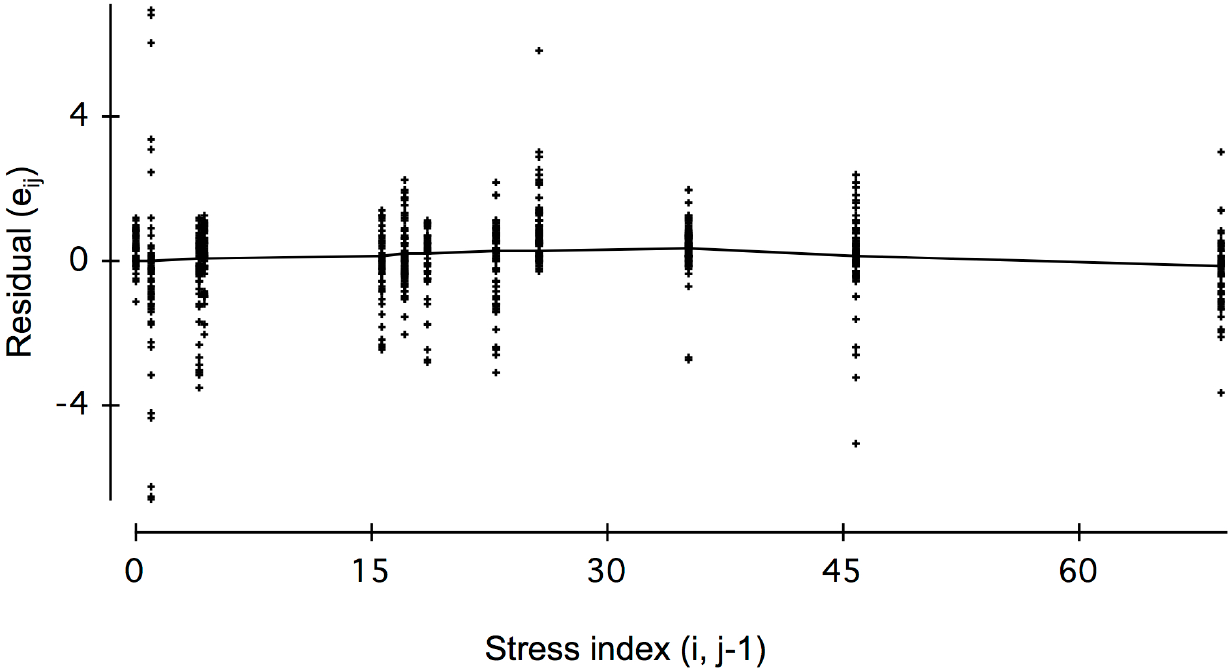
Yearly residuals of Eq. (3) in relation with the stress index of previous year, with the lowest smoothing line (DataDesk, 2011) showing the dependence of stand density on water stress

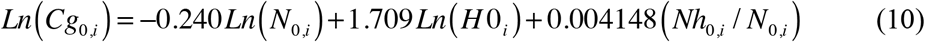

(n = 54, s = 0.100 and probability level *P* ≤ 0.0001 for each variable)

Ln(*Cg*_*0,i*_) appeared to increase linearly with the log-transformed fertility index Ln(*H0*_*i*_) and the initial proportion of beech trees (*Nh*_*0,i*_/N_0,i_), and decrease with Ln(*N*_*0,i*_).

Then, the random parameter ∂_i_ attached to each plot appeared to be a function of plot fertility (*H0*_*i*_), initial density of trees (*N*_*0,i*_) and initial proportion of beech trees (*Nh*_*0,i*_/N_0,i_) in the stand (Eq. 9 and Eq.10)

The variations of residual errors *e*_*ij*_ of Eq. 3 were also examined in relation with yearly varying stand variables, that is water stress indexes of current year *j* and of preceding year *j-1*, called respectively *Stressj* and *Stressj-1*. A slight relation was found between *e*_*ij*_ and *Stressj-1*, exhibiting a quasi-linear decrease of *e*_*ij*_, that is an increase of tree mortality above a stress level equal to 30 (Fig.10).

### 3.3 Individual tree mortality

The probability of individual tree mortality (*P*_*m*_) was analyzed by using the linear model described by Equation (5). In addition to potential individual mortality factors, possible interactions between them were also examined.

Two models presenting the highest fitting AUC values and comprising slightly different variables (Wald test, 5% significance level), more especially characterizing stand density, were compared: one included the number of trees per ha relatively to initial number (*N/N*_*0*_) and the other one, the relative density index (*RDI*). The fitting of these two models with *R* (“glm”, 2012) and its package “ROCR” compared as follows (Table 7):

**Table 7.** The two logistic models considered for tree mortality — based on Eq. 5 — with their characteristics: significant variables and fitting quality criteria (AIC and AUC)

| Model | Variables | AIC | AUC |
| --- | --- | --- | --- |
| including $N/N_0$ | C130/Cg, $N/N_0$, H0<br>Stress<sub>j-1</sub>, Stress<sub>j-1</sub>*$N/N_0$<br>Species | 5704 | 0.90 |
| including RDI | C130/Cg, $N_0$, RDI, H0,<br>Species | 6249 | 0.87 |

The first model including N/N_0_ was retained as it presented a slightly higher value for the *AUC* criterion and a lower value for the *AIC* criterion; moreover, it did not require to estimating one or more variables included in the model (like RDI for example in the second model). Thus, the retained model showed an excellent discriminatory power (area under the ROC curve equal to 0.90).

The predictive variables of Eq. 5 and the coefficients attached to each of them appear in Table 8 with their statistics: all coefficients are significant. The predictive logit function for tree mortality can then be written as follows:

**Table 8.** Parameter estimates and statistics of Equation (5) fit to the tree mortality data of the 64 plots *i* (*i* = 1 to 64) for years *j* (*j* = 2005 to 2012)

| Variables | Coefficient estimate | Std. Error | Wald Statistic* | P value |
| --- | --- | --- | --- | --- |
| Constant ($a$) | 6.673 | 0.742 | 80.96 | $\leq 0.0001$ |
| $C130_{i,j}/Cg_{i,j}$ | -6.464 | 0.211 | 940.90 | $\leq 0.0001$ |
| $N_{i,j}/N_{0,i}$ | -3.866 | 0.323 | 143.50 | $\leq 0.0001$ |
| $H0_i$ | -0.253 | 0.056 | 20.63 | $\leq 0.0001$ |
| Species |  |  |  |  |
| - beech ($Sp_b$) | -1.283 | 0.065 | 387.80 | $\leq 0.0001$ |
| - hornbeam ($Sp_h$) | -0.828 | 0.058 | 204.40 | $\leq 0.0001$ |
| - others ($Sp_o$) | 2.111 | 0.097 | 479.30 | $\leq 0.0001$ |
| $\text{Stress}_{j-1}$ | 0.093 | 0.011 | 74.57 | $\leq 0.0001$ |
| $\text{Stress}_{j-1} * N_{i,j}/N_{0,i}$ | -0.102 | 0.015 | 49.11 | $\leq 0.0001$ |

\*The *Wald* statistic, analogous to the *t*-test in linear regression, is used to assess the significance of coefficients. The Wald statistic is the ratio of the square of the regression coefficient to the square of the standard error of the coefficient and is asymptotically distributed as a chi-square distribution. It can be used carefully to assess the contribution of individual predictors.
(https://www.liquisearch.com/logistic_regression/coefficients/wald_statistic)

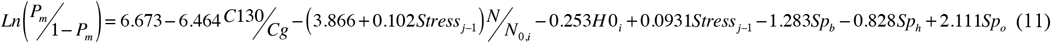

in which each “*species* indicator” (*Sp*_*b*_ for beech, *Sp*_*h*_ for hornbeam *and Sp*_*o*_ for others) in the equation takes the value 1 if the tree is of the concerned species, 0 otherwise.

## 4 Discussion

### 4.1 Size-density trajectories model

To fit the size-density trajectories of the mixed beech-based plots, the hypothesis of alignment of the inflexion points of the trajectories was made on the basis of previous studies on various species (Le Goff & al. 2021, Ningre & al. 2016a, 2016b, 2019). Moreover, the hypothesis of parallelism between mortality onset and reach of maximum density lines could be retained for some of these species including beech (Le Goff & al. 2021, Ningre & al. 2019). These assumptions could be verified again in this case of beech-based mixed stands, which state that the relative density at mortality onset (in terms of RDI) is also independent of initial tree density for these stands. The size-density trajectories of the mixed beech-based stands are thus entirely determined by the initial density of the stand N_0_.

The onset of mortality and the maximum size-density lines were compared for pure even-aged stands of beech (Ningre et al. 2019) and for mixed beech-based stands of this study, which cover a comparable range of initial densities. It appears that mortality starts “later” – that is for larger dimensions – in the mixed stands studied (Fig. 11), which means that beech can support higher densities when mixed with hornbeam, the most important species mixed with it at this stage. On the opposite, the maximum density reached by pure and mixed beech stands seems similar (Fig. 11). Then, the density at which mortality begins is higher for the mixed beech-based stands than for pure beech stands (RDI_0_ values of 0.50 and 0.29 respectively).

**Fig. 11.**
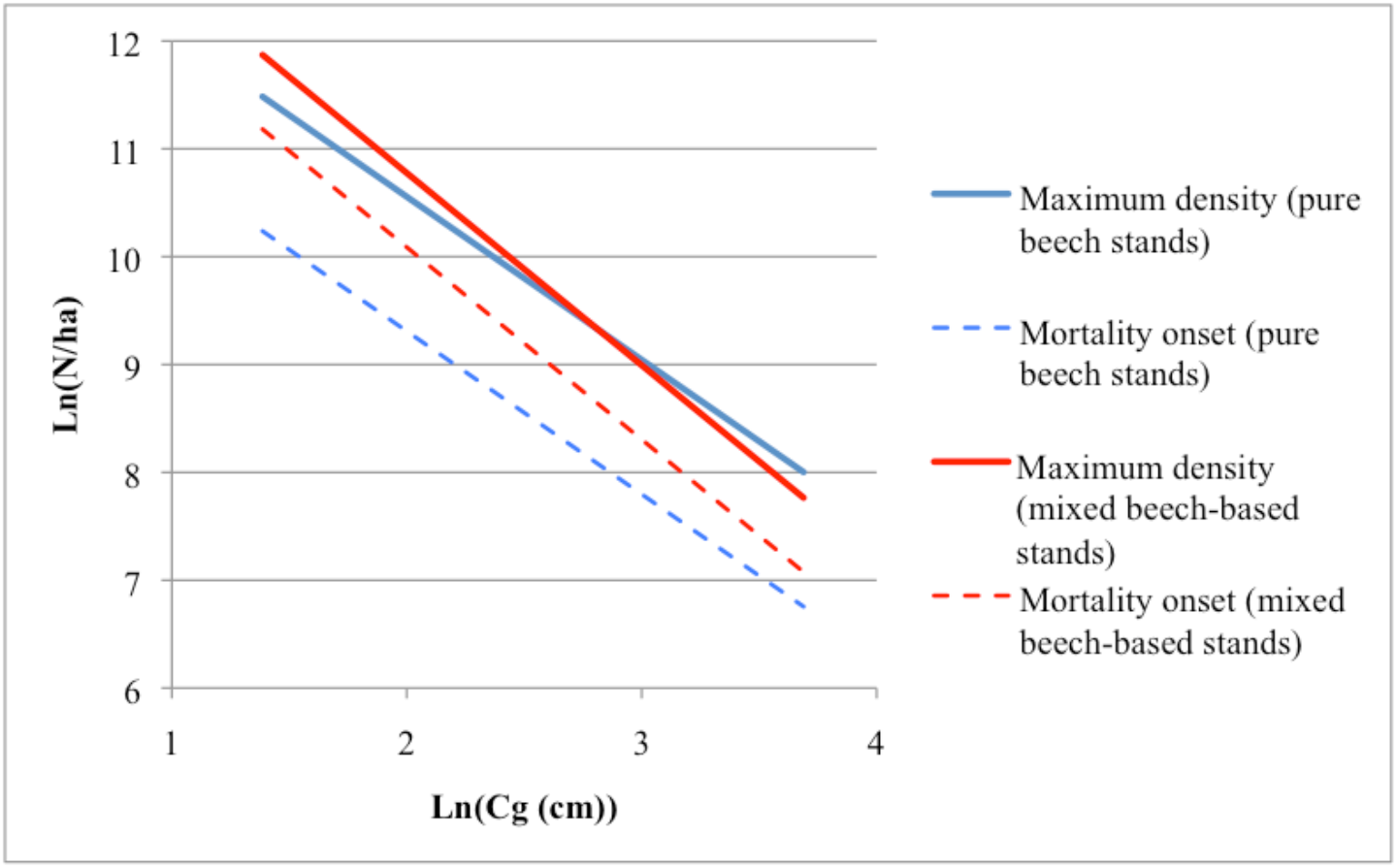
Comparison of characteristic size-density lines for mortality onset and reach of maximum density for the beech mixed stands of this study and for the pure beech stands of the density trial of Lyons-la-Forêt (Ningre et al. 2019)

The characteristics of the size-density trajectories with different initial densities (N_0_) were calculated using Eqs. (3), (7) and (8) (Table 9) and the corresponding size-density trajectories were represented, together with the common size-density lines (Fig 12). A specific characteristic of these size-density lines modeled as here, is that the relative ratio of surviving trees (N_1_/N_0_), when the stands reach the maximum density, is a constant equal to 0.5, regardless of initial stand density (Table 9 and Fig. 12). More generally, the ratio N/N_0_ is a constant for stands of a given RDI, between mortality onset (RDI_0_ = 0.5) and advent of maximum relative density (RDI_max_ = 1). RDI mortality lines could then be drawn, as it was done for oak (Ningre et al. 2019), corresponding to a given mortality rate (from 0 to 0.5). This kind of stand density management diagram may help the forester to characterize the density of mixed beech-based stands and better predict the cleaning and thinning operations to make in order to avoid mortality losses and improve tree growth.

**Table 9.** Estimations of the size-density trajectory characteristics for the range of initial densities observed, from Eqs. 3, 7 and 8: mortality onset (Cg_0_, RDI_0_) and reach of maximum density (Cg_1_, N_1_) are given for each tabulated initial density – including the mean initial density in this study (N_0_ = 18413) – and the corresponding size-density trajectories are represented in Fig. 12)

| Initial density<br>(trees per<br>ha, $N_0$) | Mortality onset | | Maximum RDI reach | | |
| --- | --- | --- | --- | --- | --- |
| | $Cg_0$ (cm) | $RDI_0$ | $Cg_1$ (cm) | $N_1$ | $N_1 / N_0$ |
| 3000 | 23.72 | 0.50 | 51.33 | 1507 | 0.50 |
| 5000 | 17.81 |  | 38.54 | 2512 |  |
| 10000 | 12.07 |  | 26.13 | 5024 |  |
| 18413 | 8.57 |  | 18.55 | 9251 |  |
| 20000 | 8.18 |  | 17.71 | 10048 |  |
| 30000 | 6.52 |  | 14.11 | 15073 |  |
| 40000 | 5.55 |  | 12.01 | 20097 |  |
| 50000 | 4.90 |  | 10.59 | 25121 |  |
| 60000 | 4.42 |  | 9.57 | 30145 |  |
| 70000 | 4.05 |  | 8.77 | 35169 |  |

**Fig. 12.**
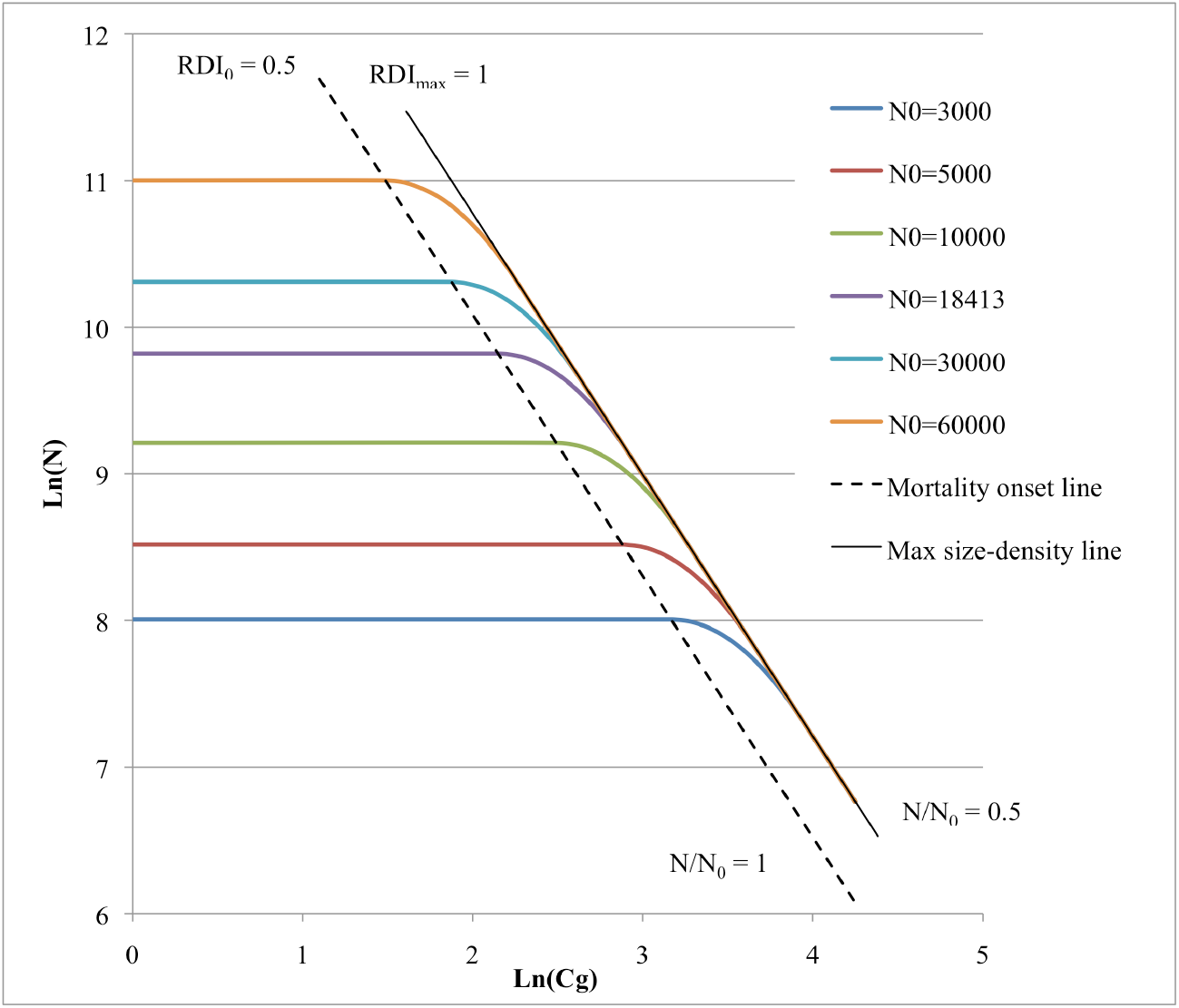
Size-density trajectories, in log-log scales, simulated for different levels of initial density (N_0_) using Eq. 3 parameters, with the lines corresponding respectively to mortality onset (RDI_0_ = 0.5 or N/N_0_ = 1) and maximum density (RDI_max_ = 1 or N/N_0_ = 0.5) where N is the total number of living trees per ha

From equations (9) and (10), it appears that the maximum density reached by the beech-based stands studied grows up slightly with fertility [*H0*], and with initial density of trees (*N*_*0*_) (Fig. 9). As *Cg*_*0*_ slightly increases also with the initial proportion of beech in the stand (Eq. 10), the maximum density seems to slightly decrease when the mixture of species increases. The same conclusions can be drawn for mortality onset which should start slightly “later” with higher levels of fertility and initial density, whereas it should slightly start “earlier” with increasing mixture of species. These findings were possible thanks to the large variability of initial stand densities and relative contribution of the main species, but also to the variation of fertility among the 64 experimental plots – H0 in 2010 varied between 11.3 m and 15.3 m – which is rarely the case in this kind of experiments.

### 4.2 Individual tree mortality model

The *logit* function for tree mortality was used to represent the effects of tree dimensions, species and climate, using its exponential form given by Eq. 6. In the chosen model including N/N_0_, as in all models with high discriminatory power, tree social status (C130/Cg) is the main explicative variable. Tree mortality increases with decreasing social status, that is with decreasing ratio C130/Cg, both for beech and for hornbeam (Fig. 13). The probabilities of mortality rise up with the progress of stand development, regardless the stress level: below a value of N/N_0_ equal to 0.5, which means that the stand has reached the maximum density (see Table 9 and Fig. 12), the probabilities of mortality are increasing. These effects appear a little more pronounced for hornbeam than for beech (Figs. 13-A/13-D; 13-B/13-E; 13-C/13-F), the probability of tree mortality being slightly higher for hornbeam for a comparable social status.

**Fig. 13.**
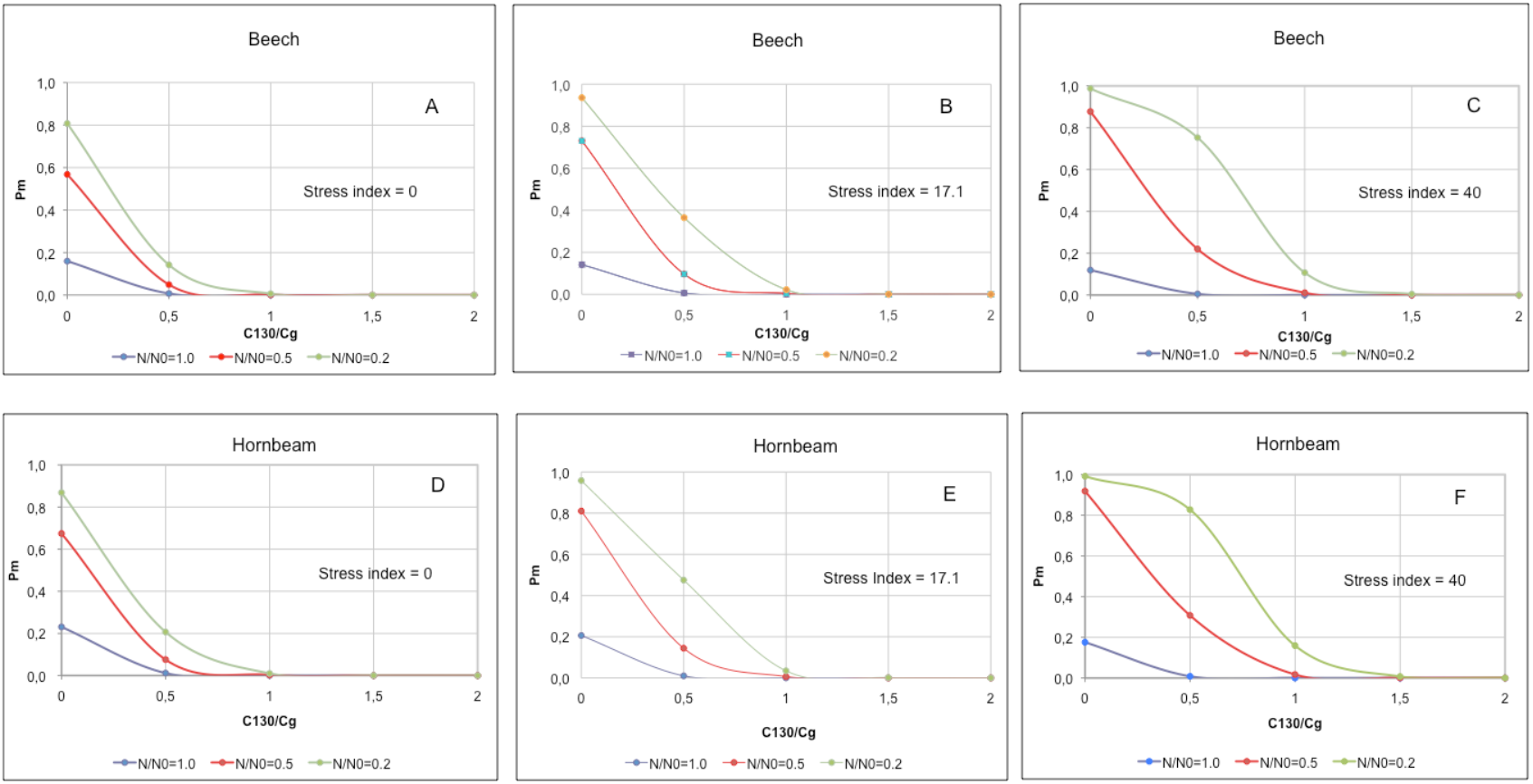
Probability of mortality of trees (*Pm*) obtained from Equation (11), respectively for beech (A, B, C) and hornbeam (D, E, F), and for varying social status of trees (C130/Cg)^8^ and for different stages of the size-density trajectories of stands measured by the ratio N/N_0_ fixed to 3 values (1, 0.5 and 0.2). Plot fertility was fixed to the mean dominant height (H0) of the 64 plots, that is 12.58 m and the stress index was fixed to 3 levels: mean stress of the period 2005-2012, that is 17.1, minimum stress (0) and maximum stress (40)

Tree mortality is also largely affected by climatic conditions, especially water deficits supported by trees, more especially during the year preceding the current one and characterized by its “stress index”. So, the probability of mortality increases for beech and hornbeam, the main species in the mixture, as the stress index climbs from 0 to 40, the interval of values observed during the study (Fig. 13-A/B/C for beech and Fig.13-D/E/F for hornbeam). Mortality affects also more and more trees of increasingly higher social status (trees of social status above 1, that of the average tree) — which is quite remarkable — in the case of the most intense stresses and for the oldest stands. Otherwise, the effect of water deficits on the probability of tree mortality appears to increase when stands are ageing (lower values of N/N_0_) and develop then at maximum density. These findings corroborate the results of ecophysiological studies conducted in the same site showing that beech (*Fagus sylvatica* L.) and hornbeam (*Carpinus betulus* L.) displayed typical characteristics of drought-sensitive species (Zapater et al. 2013). The group of other tree species, which represents only 10% of the mixture, half of them being oaks, includes species with different ecophysiological temperaments, some recognized as drought-avoidance species like *Quercus robur* and *Salix capreae* (Zapater et al. 2013), whereas the group of “other species” presents a higher probability of mortality than beech or hornbeam (see Eq. 11). Then, the other tree species present in the mixture (birch, wild cherry, aspen) could be the most drought-sensitive among the species represented in the mixture, or suffer from another growth limiting factor (space and light requirements).

### 4.3 Connection of the tree mortality model to the trajectory model

Each stand has its own mortality trajectory that depends only on N_0_ and predicts the evolution of the density of trees as a function of stand mean girth. The individual mortality model fitted, which includes N/N_0_, is therefore well suited to being combined with the size-density trajectory model for predicting at each stage of development the mortality in even-aged beech-based stands. The size-density trajectories allow the prediction of stand density (and global mortality) during stand development, given initial stand density and a growth model to predict the evolution of mean Cg (Ningre et al. 2016b), and the tree mortality model allows to predict which trees will die, given their probability of dying^9^ that depends on species, tree social status (C130/Cg), fertility conditions (H0), stand dynamics (N/N_0_) and water stress index (stress_j-1_) of the year (j-1) preceding the current one (j).

## Conclusion

This study on mixed beech-base stands is a follow-up to similar studies conducted earlier on various species growing in pure even-aged stands. The piecewise model used to represent previously the size-density trajectories of pure even-aged stands of beech, oak, Douglas-fir and ash proved its ability to represent the size-density trajectories of a mixture of beech and other hardwood species (mainly hornbeam, the major species mixed with beech in the studied regeneration).

Moreover, as for the previous species studied, natural mortality appeared here to happen at a constant relative density, independently of initial stand density. When comparing to pure beech stands in Lyons-la-Forêt (Ningre et al. 2019), the relative density at which mortality first occurred appeared higher in the mixed experimental beech stand in Hesse forest (0.5 compared to 0.29), whereas pure beech and mixed beech-based stands presented a similar maximum density. Then, beech seems to better support competition in the young stages when mixed with other hardwoods like hornbeam, maybe in relation with lower space requirements of hornbeam or with a better ability of beech to compete for space (in relation with lower light requirement).

As in other studies, the *logit* model proved here useful to describe the individual mortality of trees. The social status of trees, represented by the relative tree girth (C130/Cg) appeared as the main factor of mortality, the smaller trees with a low ratio C130/Cg being the most likely to die. Furthermore, the different species in the mixture have not an equal probability to die: beech and hornbeam seem to be the most resistant with a slight tendency towards higher mortality in the case of hornbeam. The group of “other species” appears as the less resistant, maybe in relation with higher light requirements that make them more susceptible to inter-tree competition when mixed with more shade tolerant species like beech and hornbeam. Climatic and soil conditions play also an important role through the limitations of water availability, especially those occurring the year prior to the ungoing growing season for the most drought-sensitive species in the mixture.

Finally, this study showed how size-density trajectories could be mixed with individual mortality models in tree “distant-dependent” growth simulators to predict mortality in pure or mixed even-aged stands.

## Acknowledgments

The growth and mortality data of the experiment conducted in Hesse forest (NE France) were obtained by the UMR LERFoB (Installation Expérimentale Croissance) at INRA*e* - Nancy. The experimental trial, the basis for our study, was set up in a natural beech regeneration studied for ecophysiological functioning by A. Granier, a former scientist at INRA*e*-Nancy. The data used were collected under the supervision of R. Cantá, assisted by the forest research technicians F. Bordat and G. Marechal for tree measurements and F. Vast for soil descriptions. Finally, we wish to thank André Granier and Fleur Longuetaud for their respective assistance in calculating water stress index with the model “*Biljou*” and in implementing the “ROCR” package in R.

## Annex

characteristics of the residuals of the fitted size-density trajectories equation (Eq. 3)

## Footnotes

1 Some plots were excluded from the study since year 2007, due to tree cuttings performed for the installation of ecophysiological experimental material.

2 The ht-c130 model was as follows : Ln(ht) = a + b Ln(c130) where Ln stands for the neperian logarithm

3 In that case, we hypothesize that exists a maximum size-density line independently of the stand composition or that the effect of species mixture is included in the plot effect on parameter *a*, that is 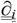

4 Core samples could not be obtained for the sample points 7, 8, 9, 10, 15 and 16, due to the presence of experimental facilities (shelter, scaffolding and trench) installed in the site

5 The water stress index includes at the same time the duration and the intensity of the soil water deficit (more details in *Biljou*, 2014)

6 Year 2000 was also considered, as the stress index of the preceding year appeared as a predictor of tree mortality for the current year (see section 3.3): the climatic data were obtained from Hesse-1.

7 A comparison of drought stress indices in beech forests is subject to a modeling study by Vilhar (2016)

7 This equation was established on a lower number of plots than equation (9) due to a lower number of values for dominant height (H0_i_) in 2010

8 Moreover, the stress index appeared linearly linked to the water holding capacity (RU) of the soil, and then the mean stress indexes of a given year for all sample plots is also the stress index of a plot presenting the mean RU of these sample plots.

8 The social status of trees on Fig. 13 was limited to the interval 0-2 as the probability of tree mortality is null above 2.

9 The onset of mortality corresponded to the year following that when the maximum number of trees was reached in a given plot and then N_0_ was not necessarily the initial number of trees of that plot (“new” trees may appear or become countable)

9 It just needs to sort trees by their probability of dying and choose the ones with the higher probabilities of dying requested to fit the size-density trajectory.

10 Cg_1_ is fully determined by Cg_0_ and N_0_ (see Ningre et al. 2016a, 2019).

11 The linearity of the relation between ln(N_0_) and Ln(Cg_0_) could be verified during the fitting process

12 Only the trees that died (*st* = 1) or stayed alive (*st* = 0) from one inventory to the next one were retained for mortality analysis; trees removed for a research purpose or just missing were not considered.

13 In Table 4 and in the text, “initial” means “before mortality appears”

6 The length of the trajectories may vary from plot to plot due to different monitoring times (see & 2.3).

